# Lentiviral vector-based SARS-CoV-2 pseudovirus enables analysis of neutralizing activity in COVID-19 convalescent plasma

**DOI:** 10.1101/2020.12.28.424590

**Authors:** Cevriye Pamukcu, Elif Celik, Ebru Zeynep Ergun, Zeynep Sena Karahan, Gozde Turkoz, Mertkaya Aras, Canan Eren, Uluhan Sili, Huseyin Bilgin, Ilke Suder, Baris Can Mandaci, Baran Dingiloglu, Ozge Tatli, Gizem Dinler Doganay, Safa Baris, Nesrin Ozoren, Tolga Sutlu

**Author notes:** Authors contributed equally (Elif Celik, Ebru Zeynep Ergun, Zeynep Sena Karahan). **Correspondence:** Dr. Tolga Sutlu, Boğaziçi University, Department of Molecular Biology and Genetics, Kuzey Kampus, Kuzey Park 320. 34342 Bebek / Istanbul / Turkey.

## Abstract

As the COVID-19 pandemic caused by Severe Acute Respiratory Syndrome Coronavirus 2 (SARS-CoV-2) continues to spread around the globe, effective vaccination protocols are under deployment. Alternatively, the use of convalescent plasma (CP) therapy relies on the transfer of the immunoglobulin repertoire of a donor that has recovered from the disease as a means of passive vaccination. While the lack of an effective antiviral treatment inadvertently increases the interest in CP products, initial clinical evaluation on COVID-19 patients revealed that critical factors determining the outcome of CP therapy need to be defined clearly if clinical efficacy is to be expected. Measurement of neutralizing activity against SARS-CoV-2 using wildtype virus presents a reliable functional assay but the availability of suitable BSL3 facilities for virus culture restricts its applicability. Instead, the use of pseudovirus particles containing elements from the SARS-CoV-2 virus is widely applied to determine the activity of CP or other neutralizing agents such as monoclonal antibodies.

In this study, we present our approach to optimize GFP-encoding lentiviral particles pseudotyped with the SARS-CoV-2 Spike and Membrane proteins for use in neutralization assays. Our results show the feasibility of pseudovirus production using a C-terminal truncated Spike protein which is greatly enhanced by the incorporation of the D614G mutation. Moreover, we report that the use of Sodium Butyrate during lentiviral vector production dramatically increases pseudovirus titers. Analysis of CP neutralizing activity against particles pseudotyped with wildtype or D614G mutant Spike protein in the presence or absence the M protein revealed differential activity in CP samples that did not necessarily correlate with the amount of anti-SARS-CoV-2 antibodies.

Our results indicate that the extent of neutralizing activity in CP samples depends on the quality rather than the quantity of the humoral immune responses and varies greatly between donors. Functional screening of neutralizing activity using pseudovirus-based neutralization assays must be accepted as a critical tool for choosing CP donors if clinical efficacy is to be maximized.

## Introduction

Severe acute respiratory syndrome coronavirus 2 (SARS-CoV-2) emerged in late 2019 in China and has rapidly affected millions of people around the world leading to the COVID-19 pandemic (1–3). While the race towards an effective and long-lasting SARS-CoV-2 vaccine is in progress, there is still no effective golden standard antiviral therapy against COVID-19. Alternatively, passive immunotherapy approaches including the use of intravenous immunoglobulins (4), transfer of convalescent plasma (5) and the use of recombinant monoclonal antibodies (mAbs) as well as mAb cocktails (6–8) are currently under clinical investigation.

Convalescent Plasma (CP), used historically to treat infectious diseases (9–11), is a passive immunotherapy approach relying on the transfer of antibodies present in the serum of a person that has recently recovered from the same infection. Rich in immunoglobulins and neutralizing antibodies (nAbs), CP may help to direct complement responses against SARS-CoV-2 particles as well as prevent viral entry by blocking receptor interaction, resulting in decreased viral replication (12–14). Mainly due to the lack of a better option, CP therapy has often been utilized as a potential option for treatment of COVID-19 patients (12, 15, 16).

Concentration of nAbs in CP is expected correlate with the effectiveness of the treatment (16, 17) and it has been suggested that there is a correlation between SARS-CoV-2 specific antibody titers in CP and the severity of COVID-19 pneumonia experienced by the donor (18–21). Since not all antibodies against SARS-CoV-2 antigens can be expected to have a neutralizing effect, it is critical to measure the neutralizing activity of CP for maximizing clinical benefit to the recipient. Evaluation of neutralizing activity can be carried out using the wildtype SARS-CoV-2 virus in a Biosafety Level 3 (BSL3) facility (22) or more commonly using a pseudovirus-based system in a BSL2 facility (14, 23)

Sequencing of the SARS-CoV-2 RNA genome has revealed numerous genetic variants in the Spike protein since the onset of the pandemic. Among these, none has attracted as much attention as the D614G mutation located just outside the Receptor Binding Domain (RBD) of the Spike protein which was shown to increase SARS-CoV-2 replication and infectivity (24–28). Viral entry of SARS-CoV-2 is facilitated by binding of its extensively glycosylated Spike protein to angiotensin-converting enzyme 2 (ACE2) located on the plasma membrane of host cells (29). RBD of Spike protein is shown to be crucial for this interaction and is identified to be a common target for neutralizing antibodies (30). Therefore, pseudovirions that express either full length S protein (31) or its RBD are used in neutralization assays (30, 32). Commonly used platforms for pseudovirus generation rely on Vesicular Stomatitis Virus (VSV)-based or Human Immunodeficiency Virus 1 (HIV-1)-based vector systems (23, 33, 34). A number of studies have also included the Membrane (M), nucleocapsid (N) and envelope (E) proteins of SARS-CoV-2 for improved virus-like particle (VLP) production (35, 36). In line with this, Xu et al. (37) reported that M and E proteins were required for efficient pseudovirus formation.

In this study, we present our efforts on developing a lentiviral vector-based pseudovirus system for SARS-CoV-2 neutralization assays. We show that the use of a D614G mutant Spike sequence and/or the use of the histone deacetylase inhibitor Sodium Butyrate significantly increased pseudovirus production while the inclusion of M protein did not seem to further enhance production. Additionally, no difference was observed in the susceptibility of pseudovirus particles carrying the D614G mutation or M protein to neutralization by soluble ACE2-IgG, confirming Spike-mediated entry of pseudovirus into target 293FT cells engineered to overexpress the hACE2 gene. Analysis of CP samples using this assay revealed that a significant number of CP donations have very low neutralizing activity. This indicates that neutralization assays such as the one presented in this study should be routinely used to analyze CP activity in order to ensure clinical benefit for patients receiving therapy.

## Materials and Methods

### Cell lines

The 293FT cell line was purchased from Thermo Fisher Scientific (MA, USA) and was maintained in DMEM (LONZA, Switzerland) supplemented with 10% fetal bovine serum (FBS) (HyClone, GE Healthcare, USA), 0.1 mM nonessential amino acids (HyClone), 6 mM L‐glutamine (HyClone), 1 mM sodium pyruvate (HyClone), and 20 mM HEPES (HyClone). Cells were passaged 1:4 to 1:6 depending on confluency, never allowed to reach above 90% and were used up to 20 passages. Genetically modified 293FT-hACE2 cells were selected with a final concentration of 1 mg/ml Puromycin (Sigma-Aldrich, USA).

### Plasmids

The following plasmids were used for the study: pMDLg/pRRE and pRSV-Rev packaging plasmids were gifts from Didier Trono (Addgene plasmid #12251 and #12253). pCMV-VSV-G plasmid encoding for the vesicular stomatitis virus glycoprotein was a gift from Bob Weinberg (Addgene plasmid #8454). LeGO-G2, LeGO-iT2 and LeGO-iT2puro plasmids were gifts from Boris Fehse (Addgene plasmids #25917 and #27343). pcDNA3.1+_C-(K)DYK-ACE2 (NM_021804.2, OHu20260) encoding for *Homo sapiens* angiotensin I converting enzyme 2 (hACE2) was a gift from Şaban Tekin (TUBITAK-MAM). pTwist EF1 Alpha-SARS-Cov-2-S-2xStrep plasmid encoding for the SARS-CoV-2 Spike protein was a gift from Nevan Krogan (Addgene plasmid #141382). pcDNA3-sACE2(WT)-Fc(IgG1) and pcDNA3-SARS-CoV-2-S-RBD-sfGFP plasmids were gifts from Erik Procko (Addgene plasmids #145163 and #141184). pGBW-m4134096 encoding for SARS-CoV-2 M protein was a gift from Ginkgo Bioworks (Addgene plasmid #152039).

### Cloning

For the overexpression of hACE2 protein on 293FT cells, LeGo-ACE2-iT2 and LeGo-ACE2-iT2puro vectors were constructed. All primers were synthesized by Sentromer DNA Technologies, TR. hACE2 region from pcDNA3.1+_C-(K)DYK-ACE2 was Polymerase Chain Reaction (PCR) amplified using forward primer containing NotI Cut site, 5’-AAT <u>GCG GCC GCC</u> ACC ATG TCA AG-3’ and the reverse primer containing NotI Cut Site 5’-TAT <u>GCG GCC GCT</u> TAA AAG GAG GTC TGA A-3’ (restriction sites underlined). For preparation of the vector, LeGO-iT2 and LeGO-iT2puro plasmids were cut by NotI-HF (New England Biolabs (NEB), USA) and ligation with the NotI digested PCR products was performed for 1 hour at room temperature with T4 DNA ligase (NEB). Colonies were screened by restriction digestion for the directional insertion of PCR products and the resulting lentiviral vectors were validated by Sanger sequencing.

Full-length SARS-CoV-2 Spike gene and the 19 aminoacid truncated version without the endoplasmic reticulum retention signal (ERRS) (SpikeΔ19) regions from pTwist-EF1-Alpha-SARS-Cov-2-S-2xStrep were PCR amplified using the same forward primer containing BamHI cut site, 5’-TTA T<u>GG ATC C</u>GC CGC CAC CAT GTT TGT T-3’ and different reverse primers containing BamHI cut sites 5’-GGC G<u>CG GAT CC</u>T TAC GTG TAG TGC AAT T-3’ and 5’-GTG T<u>GG GAT CC</u>T TAG CAG CAA CTA CCG C-3’. pCMV-VSV-G was cut by BamHI-HF (NEB) for removing the VSV-G coding region and the digested PCR products were cloned into this site. Ligations were performed with T4 DNA ligase (NEB) for 15 minutes at room temperature followed by 1 hour at 16°C. The generated vectors were named pCMV-Spike and pCMV-SpikeΔ19 and constructs were validated by Sanger sequencing.

The D614G variants on both pCMV-Spike and pCMV-SpikeΔ19 plasmids were generated by site-directed mutagenesis. For point mutation on D614, pCMV-Spike and pCMV- SpikeΔ19 plasmids were PCR amplified using a forward primer 5’-CAG TTC TTT ATC AGG GCG TGA ATT GTA CAG AG-3’ and a reverse primer 5’-TCT GTA CAA TTC ACG CCC TGA TAA AGA ACT GC-3. DpnI (NEB) restriction enzyme was used to remove unmodified plasmids and the PCR products were transformed into Top10 (Thermo Scientific, USA) *E.coli* for amplification. The generated vectors were named pCMV-Spike(D614G) and pCMV-SpikeΔ19(D614G) and the mutagenesis was validated by Sanger sequencing.

### Production of recombinant soluble proteins

For production of soluble proteins (RBD-GFP and ACE2-IgG1), 293FT cells (6×10^6^) were seeded in poly-L-lysine (Sigma, USA) coated 100mm tissue culture dishes (Corning, USA). When cells reached 70% confluency on the following day, they were transfected with the 15 μg of the plasmids (either pcDNA3-SARS-CoV-2-S-RBD-sfGFP or pcDNA3-sACE2(WT)-Fc) using calcium phosphate precipitation in the presence of 25 μM chloroquine. After 8-10 h incubation, medium was changed with complete DMEM containing 5 mM sodium butyrate (NaBut). The supernatants were collected at 48-60 h after medium change and centrifuged at 300g for 5 mins to remove cell debris. After centrifugation, the supernatants were filtered through 0.45μm filters. Filtered supernatants of RBD-GFP were aliquoted and stored at −20°C as required.

Filtered supernatant of ACE2-IgG1 was diluted 1:3 with Buffer A (20 mM Sodium Phosphate, pH 7.4, 150 mM NaCl) and captured by HiTrap MabSelectSure (Cytiva) 1mL column using ÄKTA Avant 25 (GE Healthcare). The column was then washed with 7.5 CV Buffer A. Captured ACE2-IgG1 was eluted with 10 CV Buffer B (100 mM Glycine, pH 3.0) and fractionated in tubes already containing 15 μL Buffer C (1M Tris pH 8.5). Collected proteins were buffer exchanged into PBS via multiple rounds of centrifugation in 10 kDA ultrafiltration tubes (Merck, USA). The purified protein content was measured by Bradford assay (B6916, Sigma-Aldrich) and analyzed under reducing conditions with 12% SDS-PAGE, followed by wet transfer on PVDF Membrane (Merck). The membrane was blocked with 5% skimmed milk in TBS-T and incubated overnight at 4°C with polyclonal rabbit anti human-ACE2 antibody (#4355), (Cell Signaling Technology (CST), USA). The dilution was prepared as 1:1000 in %5 Bovine Serum Albumin (BSA) in TBS-T. After washing with TBS-T, membrane was treated with HRP-conjugated anti-rabbit IgG (#7074, CST) and incubated 1h at room temperature. The band was visualized using ECL (Advansta, USA) on a gel documentation system (Syngene, G BOX) for validation of the purity and size of the produced ACE2-IgG protein.

### Generation human ACE2 over-expressing cells

For production the LeGo-hACE2-iT2puro vector, 293FT cells (6×10^6^) were seeded in a poly-L-lysine coated 100mm tissue culture dish and when cells reached %70 confluency on the following day, they were transfected with the following mixture of plasmids; 7.5μg of LeGo-hACE2-iT2puro, 3.75μg of pMDLg/pRRE, 2.25μg of pRSV-Rev and 1.5μg of phCMV-VSV-G using Calcium Phosphate transfection in the presence of 25 μM chloroquine. After 8-10h medium was changed and the supernatants were collected at 24h after medium change, filtered with 0.45μm filters, mixed at a 1:5 (v/v) ratio with 50% PEG8000 solution and stored at 4°C overnight. Next day, lentiviral particles were concentrated by centrifugation at 3000g for 30 minutes. Supernatants were removed and pellets resuspended in 500 μl serum-free DMEM.

For generation of 293FT-hACE2 cells, 1×10^6^ 293FT cells were seeded into a T25 flask one day prior to viral transduction. Cells were transduced with the concentrated LeGo-ACE2-iT2puro (MOI=1) vector in the presence of 8 μg/ml protamine sulfate overnight. Virus containing supernatant was completely removed the next day and cells were cultured in their regular growth media for 72 hours before tdTomato expression was assessed by flow cytometry. After puromycin selection, 293FT cells stably expressing wild type ACE2 protein were confirmed with flow cytometry.

### Flow Cytometry

For surface staining of 293FT-hACE2 cells, 2.5×10^5^ cells were washed once with PBS and incubated at RT for 1 hour with 300 μl of RBD-GFP supernatant collected as described above. The stained cells were washed with PBS and data acquisition was performed on BD Accuri C6 (Becton, Dickinson and Company, USA).

For data acquisition of neutralization assays, 25μ of Trypsin-EDTA (0.05%) with phenol red (HyClone) was added for each well on the 96-well plate and incubated at 37°C for 5 min. Cells were collected from wells by pipetting with PBS (containing 2%FBS and 2mM EDTA) and data acquisition was performed on a BD Accuri C6 (Becton, Dickinson and Company, USA). All flow cytometry data were analyzed with FlowJo v10.1 (BD Biosciences).

### Microscopy

For microscopy, 10^5^ 293FT-hACE2 cells were seeded in a poly-L-lysine coated coverslip placed in a 24-well plate and incubated overnight. The following day, coverslips were washed once with PBS, fixed with 250 μl of 4% Paraformaldehyde (PFA) for 10 minutes at 37°C and were washed two more times with PBS. 250 μl of filtered permeabilization buffer (0.2%(w/v) Saponin, 2%(w/v) BSA) was added on the cover slips and incubated at RT for 1 hour. The permeabilized cells were washed once with PBS followed by incubation with 300 μl of RBD-GFP supernatant at RT for 1 hour. Finally, coverslips were washed twice with PBS and mounted on slides for imaging. Images acquisition was performed on a Leica TCS SP5 confocal microscope (Leica Microsystems GmbH, Wetzlar, Germany). All samples were captured with 2048*2048-pixel resolution, 40X magnification, and z-stack images of 0.6 μm. Laser power, gain, and offset were kept constant for all images. Analysis of the captured images was made by Fiji distribution of Image J software 1.52p (38). Colocalization was documented by merging the three channels.

### Pseudovirus production and titration

For production of Spike-pseudotyped lentiviral particles, 293FT cells (6×10^6^) were seeded in a poly-L-lysine coated 100mm tissue culture dish and when cells reached %70 confluency on the following day, 7.5 μg vector plasmid containing GFP reporter gene (LeGo-G2) was co-transfected using Calcium-Phosphate precipitation as described above along with 3.75 μg of pMDLg/pRRE, 2.25 μg of pRSV-Rev and indicated amounts of one of the envelope plasmids encoding the SARS-CoV-2 Spike protein (pCMV-Spike for full length wildtype Spike protein, pCMV-SpikeΔ19 for wildtype Spike protein lacking 19 aminoacids in its C terminal tail or pCMV-SpikeΔ19(D614G) for Spike protein carrying the D614G mutation and lacking the 19 amino acids in its C terminal tail). In indicated experiments 1 μg of the pGBW-m4134357 plasmid encoding for the SARS-CoV-2 M protein was included in this mixture. After 8-10h medium was changed and the supernatants containing pseudovirus particles were collected at 40-48 hours after chaning medium and filtered through a 0.45 μm syringe filter, divided into aliquots and stored at −80°C.

Virus titer was estimated by transducing 5×10^5^ 293FT-hACE2 cells per well in a 24-well plate with 250 μl viral supernatant for 16 hours in the presence of 8 μg/ml Protamine Sulfate. The transduced cells were cultured for 72 hours before GFP expression was analyzed by flow cytometry and the percentage of cells transduced was used to calculate as infectious units/per ml for the pseudovirus supernatant.

### Pseudovirus-based neutralization assay

Sixteen hours prior to infection, 1×10^4^ 293FT-hACE2 cells were seeded in 100 μl full growth media into flat bottom 96-well plates. Plasma samples were heat-inactivated in microcentrifuge tubes by heating to 56°C for 30 minutes, starting approximately 1 hour before infection of cells. At the end of heat inactivation, tubes were centrifuged in a tabletop microcentrifuge for 10 seconds at top speed to get rid of any precipitates and the supernatant was used. We used an initial plasma dilution of 1:20 and did 2-fold serial dilutions in serum-free DMEM up to 1:20480. For ACE2-IgG we used an initial concentration of 20μg/ml and did 2-fold serial dilutions in serum-free DMEM up to 19ng/ml. The plasma and/or ACE2-IgG1 dilutions were mixed with pseudovirus supernatants in triplicates in the wells of a fresh 96-well plate and incubated at 37 °C for 1 hour. At the end of the incubation these mixtures were transferred to wells with seeded 293FT-hACE2 and incubated overnight in the presence of 8 μg/ml Protamine Sulfate. Next day (14-16h post transduction), medium was changed and cells were cultured for 72 hours before GFP expression was analyzed by flow cytometry.

Results of neutralization assays were plotted by normalization to samples where no plasma was used and the half-maximal inhibitory concentration (IC50) was calculated using 4-parameter non-linear regression. The results of regression analysis were only deemed acceptable when the R^2^ value was above 0.9 and the dilution factor corresponding to the calculated IC50 was used as Neutralizing Titer 50 (NT50) values.

### Donors and convalescent plasma samples

This study was approved by The Turkish Ministry of Health’s Scientific Research Platform (05.05.2020) and by Marmara University Clinical Research Ethics Committee (08.05.2020/554) and by Boğaziçi University Institutional Review Board for Research with Human Subjects (FMINAREK) (04.08.2020-2020/07). All donors applied voluntarily to the blood bank of Marmara University Pendik Education and Research Hospital for plasma donation. They were screened and fulfilled the criteria for convalescent plasma donation set forth in the Turkish Ministry of Health’s Handbook on COVID-19 Immune Plasma Procurement and Use. All donors signed informed consent for the study. Demographic information, comorbidities, symptoms, signs, imaging and PCR results were retrospectively collected from electronic health records. Severity of the patients at the time of diagnosis was defined using the World Health Organization (WHO) ordinal scale for clinical improvement. The overall characteristics of donors are given in Table 1.

**Table 1:**
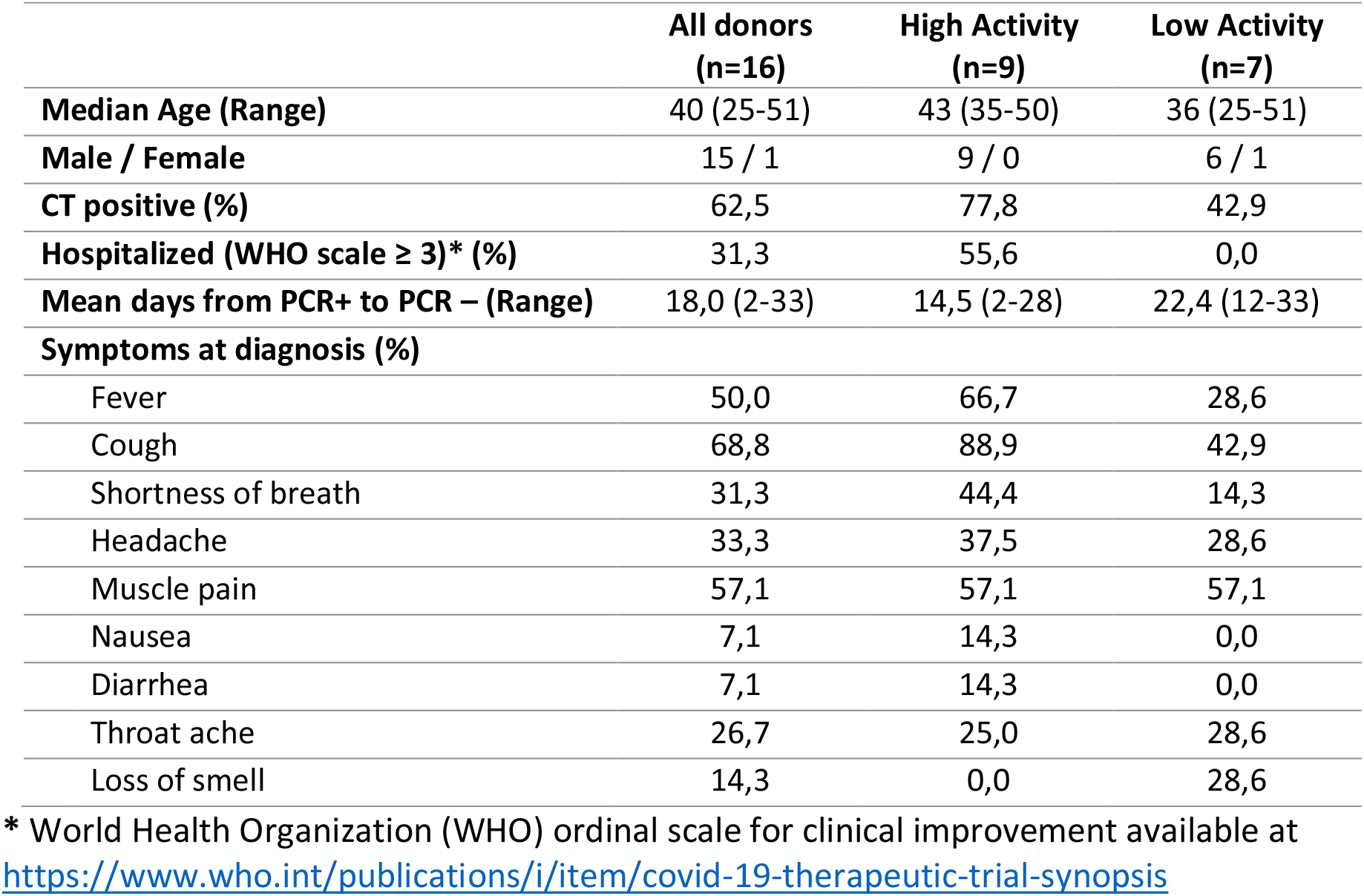
Characteristics and COVID-19 history of CP donors used in the study.

Plasma were collected from 16 convalescent donors from May to July 2020. Four healthy apheresis donors for thrombocyte collection provided control (non-convalescent) plasma samples. Criteria to become a convalescent plasma donor involved: COVID-19 infection confirmed with a positive PCR result, 14 days past after recovery with two negative PCR results or 28 days past after recovery, and a positive SARS-CoV-2 IgG antibody result. A total of 430 mL convalescent plasma was collected using apheresis method in a Haemonetics MCS+ system. Plasma were cryopreserved as 200 ml per bag, while 30 ml was spared for experimental analysis.

### Data analysis and statistics

Flow Cytometry data was analyzed using FlowJo v10.1 software (BD Biosciences). For the preparation of graphs and for statistical analysis, Prism v8.4.3 software (GraphPad Software) was used. Additional information regarding the applied statistical tests are provided in relevant figure legends.

## Results and Discussion

### Overexpression of hACE2 in 293FT cells

293FT cells were genetically modified to overexpress the hACE2 receptor in order to make them permissive to infection by Spike-pseudotyped lentiviral vectors. For this purpose, the hACE2 gene was cloned into the LeGO-iT2puro vector that codes for a bicistronic transcript harboring an IRES sequence followed by the tdTomato fluorescent protein fused with puromycin resistance gene under the control of an SFFV promoter (**Fig. 1**). This facilitates the enrichment of genetically modified cells using puromycin selection while enabling the visual tracking of hACE2+ cells by red fluorescence. After genetic modification and puromycin selection, the enrichment of tdTomato expressing cells was validated by flow cytometry. Furthermore, we investigated the surface expression of hACE2 by using RBD-GFP fusion protein in flow cytometry and confocal microscopy. Flow cytometry staining with RBD-GFP confirmed that 293FT cells modified with LeGO-ACE2-iT2puro vector expressed high amounts of the hACE2 receptor on their cell surface while control cells modified with an empty LeGO-iT2puro vector showed no signs of RBD-GFP staining (**Fig. 1A and 1B**). This was further confirmed by microscopy analysis where the localization of RBD-GFP staining in the cell surface of modified 293FT cells ensured proper expression and membrane transport of the hACE2 receptor (**Fig. 1C**).

**Figure 1:**
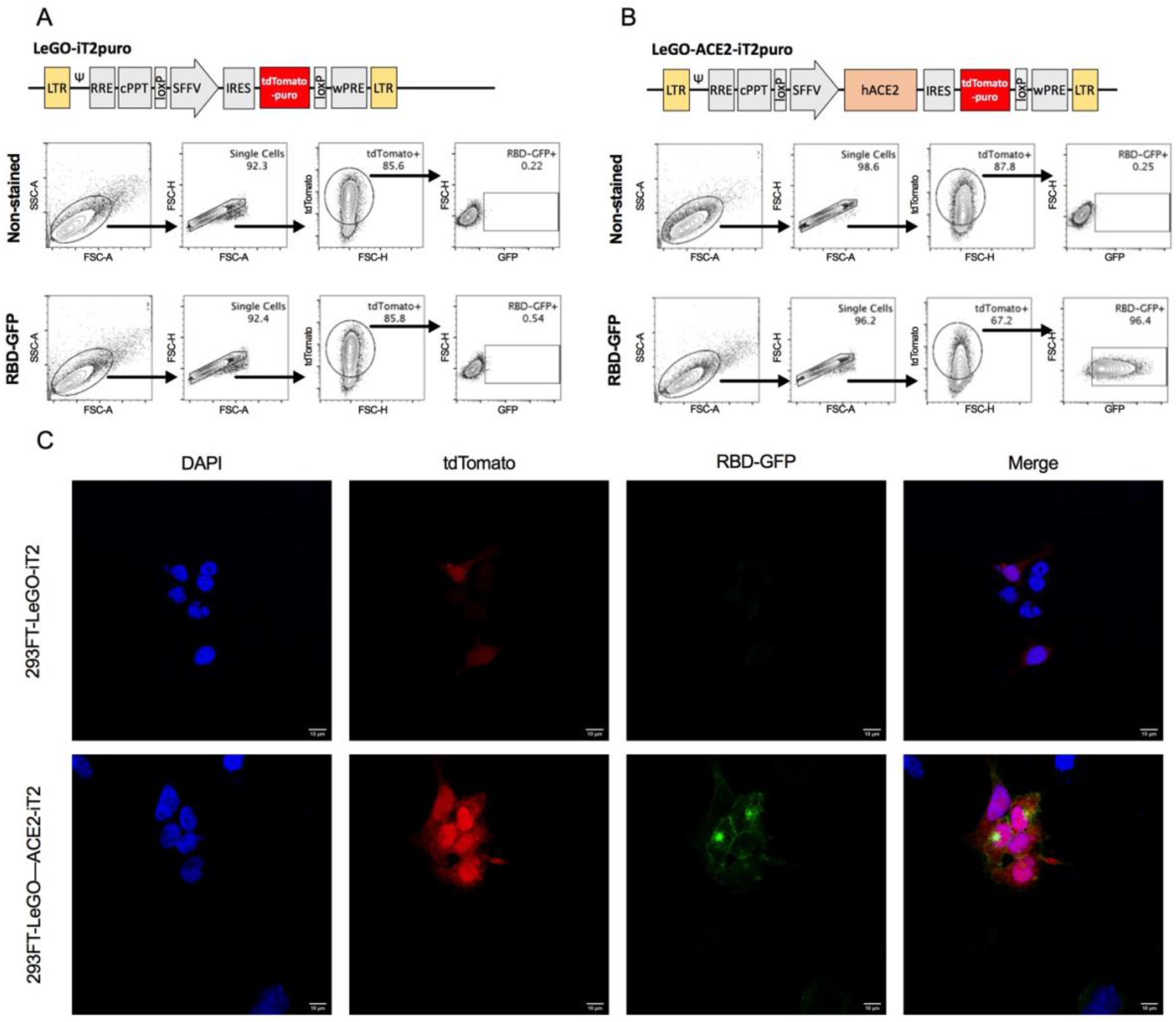
Overexpression of hACE2 in 293FT cells. Upper panels in **(A)** and **(B)** represent the maps for the empty LeGO-iT2puro vector backbone and LeGO-hACE2-iT2puro vector used to genetically modify 293FT cells. Lower panels show representative flow cytometry analysis of RBD-GFP fusion protein staining on tdTomato or hACE2-tdTomato expressing 293FT cells. **(C)** Representative confocal microscopy images generated from RBD-GFP fusion protein stained tdTomato and hACE2-tdTomato expressing HEK293FT cells.

### Production of basic Spike-pseudotyped lentiviral vector

For production of Spike-pseudotyped lentiviral vectors, SARS-CoV-2 Spike gene was cloned into the pCMV backbone used regularly for expression of the VSV-g envelope protein. The constructs prepared for pseudotyping included pCMV-Spike encoding for the wildtype Spike protein and pCMV-SpikeΔ19 encoding for the wildtype Spike protein without the last 19 amino acids that act as an endoplasmic reticulum retention signal (ERRS). Moreover, site-directed mutagenesis was employed on these two constructs to introduce the D614G mutation, resulting in two more plasmids: pCMV-Spike(D614G) and pCMV-SpikeΔ19(D614G) (**Fig. 2**). Pseudovirus production was carried out as outlined in **Fig. 3A** where one of the Spike-encoding plasmids was co-transfected into 293FT cells using calcium-phosphate precipitation along with the LeGO-G2 lentiviral transfer vector encoding the green fluorescent protein (GFP) as well as the packaging plasmids encoding for Rev and Gag/Pol. The supernatants containing pseudovirus particles were collected 48h post-transfection and tested on 293FT-hACE2 cells for infectious capacity. Analysis of GFP gene delivery by the pseudovirus into 293FT-hACE2 cells was carried out 3-days post transduction by flow cytometry (**Fig. 3E**).

**Figure 2:**
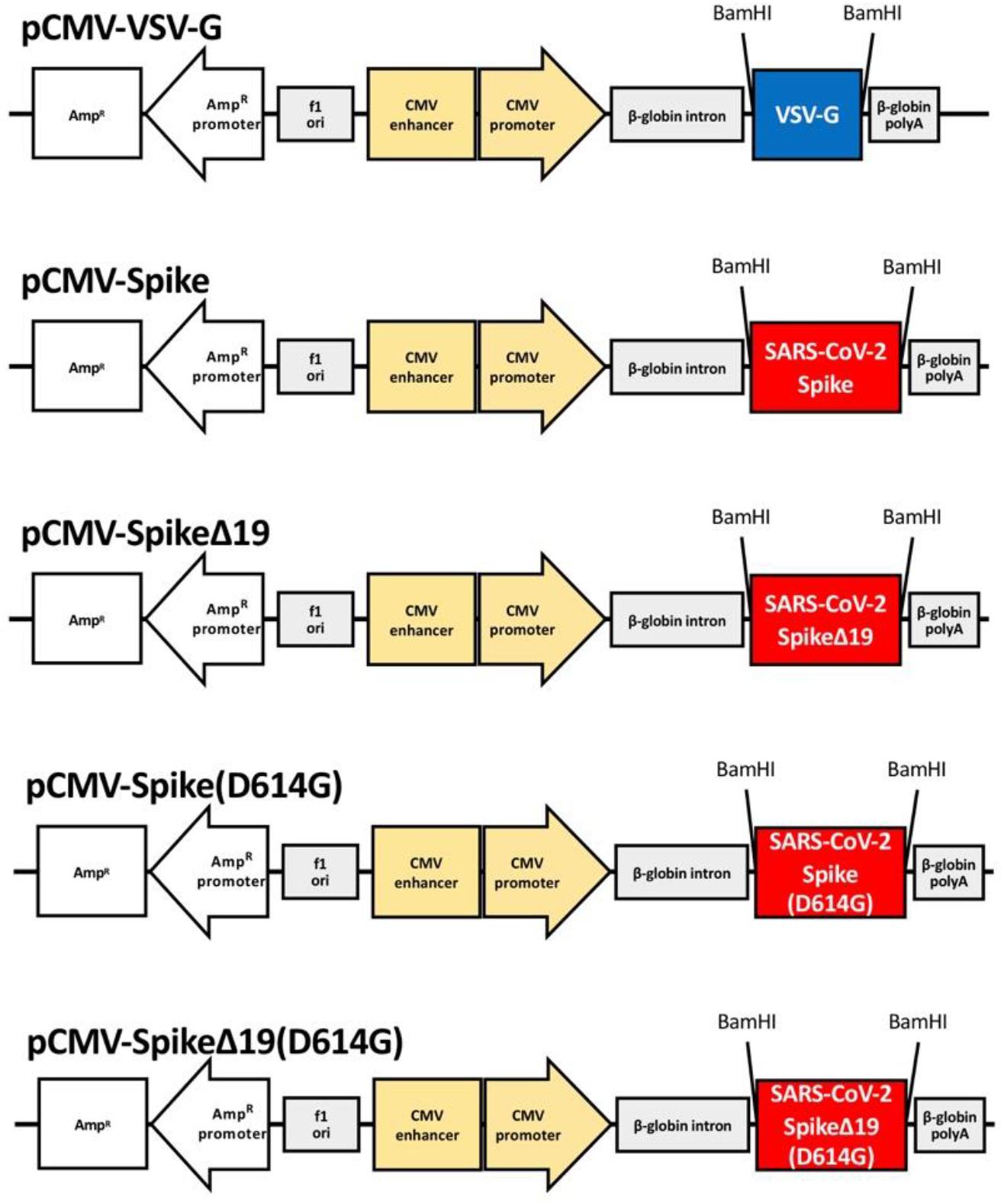
Maps of plasmids used for expression of SARS-CoV-2 Spike protein. pCMV-VSV-G backbone was used to clone different Spike sequences to be used for pseudotyping. Spike: Full length Spike protein. SpikeΔ19: Spike protein lacking 19 amino acids at the C terminal. Spike(D614G): D614G mutated full-length Spike. SpikeΔ19(D614G): D614G mutated Spike protein lacking 19 amino acids at the C terminal.

**Figure 3:**
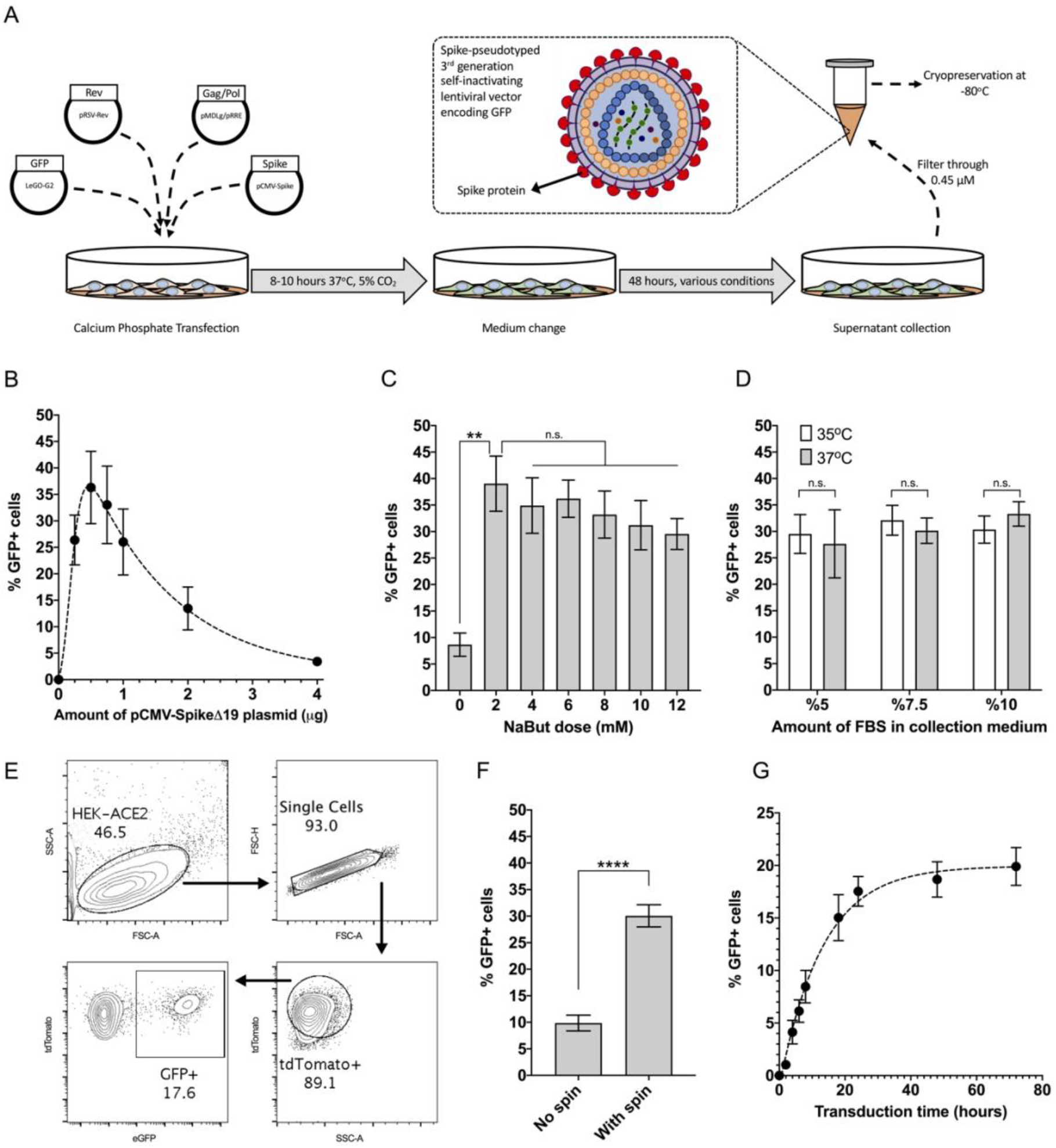
Optimization of conditions for basic pseudovirus production and transduction. **(A)** Representative protocol for the production of Spike-pseudotyped lentiviral vectors encoding GFP. Lego-G2 lentiviral plasmid was co-transfected into 293FT cells along with packaging constructs and the plasmid for Spike expression. Collected supernatants containing pseudovirus particles were used to infect 293FT-hACE2 cells. **(B)** Percentage of GFP+ 293FT-hACE2 cells after transduction with pseudovirus containing supernatants prepared with varying levels of the plasmid pCMV-SpikeΔ19 during transfection. (Data from one representative experiment runt in triplicates, Mean +/− SEM plotted) **(C)** Percentage of GFP+ 293FT-hACE2 cells after transduction with pseudovirus containing supernatants collected in cull growth medium containing different concentrations of Sodium Butyrate (Data from one representative experiment runt in triplicates. **p<0.005, n.s. not significant, One-way ANOVA, Tukey’s test) **(D)** Percentage of GFP+ 293FT-hACE2 cells after transduction with pseudovirus containing supernatants collected under different concentrations of FBS and temperatures. (Data from one representative experiment runt in triplicates. n.s. not significant) **(E)** Representative flow cytometry analysis of GFP expression 3 days after pseudovirus treatment of 293FT-hACE2 cells. Effects of **(F)** spinoculation and **(G)** exposure time to pseudovirus during transduction of 293FT-hACE2 cells. (Data from two experiments with two different batches of pseudovirus run in triplicates, **** p<0.001, paired t-test, two-tailed)

Our initial observations showed that the use of the full-length Spike gene without removal of the ERRS was ineffective for producing pseudotyped lentiviral particles possibly due to poor membrane translocation of the Spike protein. However, the use of pCMV-SpikeΔ19 plasmid successfully yielded lentiviral particles that had the capacity to infect 293FT-hACE2 cells. Our results are in line with previous reports where the removal of the ERRS sequence has been a commonly used approach for Spike-pseudotyped vector generation (33, 39) while several other studies have also reported efficient pseudovirus generation with full-length Spike sequences (40, 41). Taken together, these results indicate that it might be possible to generate low-titer pseudovirus particles using the full-length Spike protein and use it for neutralization assays after concentrating supernatants but even in these cases, the deletion of ERRS has been shown to dramatically increase titers. In our hands, the use of SpikeΔ19 for pseudovirus packaging was efficient enough to render any need for supernatant concentration obsolete.

In an effort to optimize the production of pseudovirus particles, different conditions during transfection and supernatant collection were also tested (**Fig. 3**). Initially, the amount of envelope plasmid encoding for the Spike protein within the transfection mixture was carefully titrated from 0.25 μg to 4 μg (**Fig. 3B**). While very low amounts of pCMV-Spike plasmid was not sufficient for optimal pseudovirus production, amounts higher than 1 μg presented with even lower titers due to the toxic effects of Spike overexpression by 293FT cells. We show that under these conditions, the use of 0.5 μg pCMV-SpikeΔ19 was optimal for production of Spike-pseudotyped lentiviral vector particles.

Next, we investigated the use of Sodium Butyrate (NaBut) to increase pseudovirus titers as has been repeatedly reported for various other pseudotypes in the literature (42, 43). NaBut is known to increase retroviral and lentiviral vector production capacity of 293FT cells by activating CMV enhancer and HIV-1 LTR promoter function (44). Indeed, the addition of NaBut post-transfection dramatically increased the production of Spike-pseudotyped lentiviral particles (**Fig. 3C**). While at higher doses NaBut seemed to have a toxic effect on the producer cells, we observed that the use of a 2 mM concentration was tolerable and dramatically increased pseudovirus production.

The reduction of the amount of FBS used during vector production may positively affect final titers because the stability of viral particles in serum-containing medium might be lower. However, this approach is also considered to be a “double-edged sword” as such reduction in FBS amounts may also negatively affect the viability of the producer cells, thereby decreasing the final titer (45). Similarly, lowered incubation temperatures during vector production can also increase titer due to enhanced stability of viral particles (46) or decrease the titer due to negative metabolic effects on the producer cells (47). Therefore, we carried out a set of experiments in order to investigate whether the amount of FBS used in the collection medium or the incubation temperature had any effect on pseudovirus production (**Fig. 3D**). The use of different FBS amounts (2.5-10 %) or different incubation temperatures (35°C vs 37°C) during production showed no significant effect on pseudovirus titers. Following these results, we decided to keep the standard protocol of using 10% FBS along with incubation at 37°C for pseudovirus production.

Finally, we aimed to optimize conditions for pseudovirus transduction by identifying whether the use spinoculation or long incubation times had a significant effect on the gene delivery efficiency to 293FT-hACE2 cells. Our results showed that centrifugation at 1000xg for 60 minutes during transduction yielded a significantly increased amount of GFP positive cells, especially in experiments where lower titer supernatants were used (**Fig. 3F**). We also observed that longer co-incubation of 293FT-hACE2 cells with the pseudovirus particles enhanced gene delivery efficiency up to the 24h timepoint, after which there was no further increase in the amount of GFP+ cells (**Fig. 3G**). Taking into account practicality considerations and unnecessary risk of aerosol production during spinoculation, we have opted for not utilizing spinoculation any further for the development of neutralization assays but rather relied on 24h co-incubation of the pseudovirus with 293FT-hACE2 cells.

### Production of enhanced pseudovirus

We next attempted the integration of D614G mutation into the Spike gene sequence as well as the incorporation of SARS-CoV-2 M protein into the pseudovirus particles in order to identify the effects of these modifications on pseudovirus production, stability and neutralization response. For this purpose, pseudovirus particles carrying the D614G mutation and/or M protein were initially used to infect 293FT-hACE2 cells immediately after fresh collection of the supernatant as well as after a freeze/thaw cycle at −80°C or incubation at 37°C.

Our results clearly show that the D614G mutation significantly increases the efficiency of pseudovirus production (**Fig. 4A**) and this can, at least in part, be attributed to the radically increased stability of viral particles carrying the D614G mutation which was mostly evident at 37°C where the half-life of the pseudovirus particles were increased to 12 hours compared to the 5 hours observed with the wildtype Spike protein (**Fig. 4B**). These results are in line with reports showing increased stability and *in vitro* infectivity of the D614G mutant (48–50). We also observed that the incorporation of M protein into the particles does not affect the production efficiency but might be slightly decreasing the stability of the pseudovirus. This slight decrease of stability was most evident in freeze/thaw experiments within the context of SpikeΔ19 as well as the stability of the particles at 37°C but was not significant when the more stable D614G mutant was used for pseudovirus production (**Fig. 4A and 4B**).

**Figure 4:**
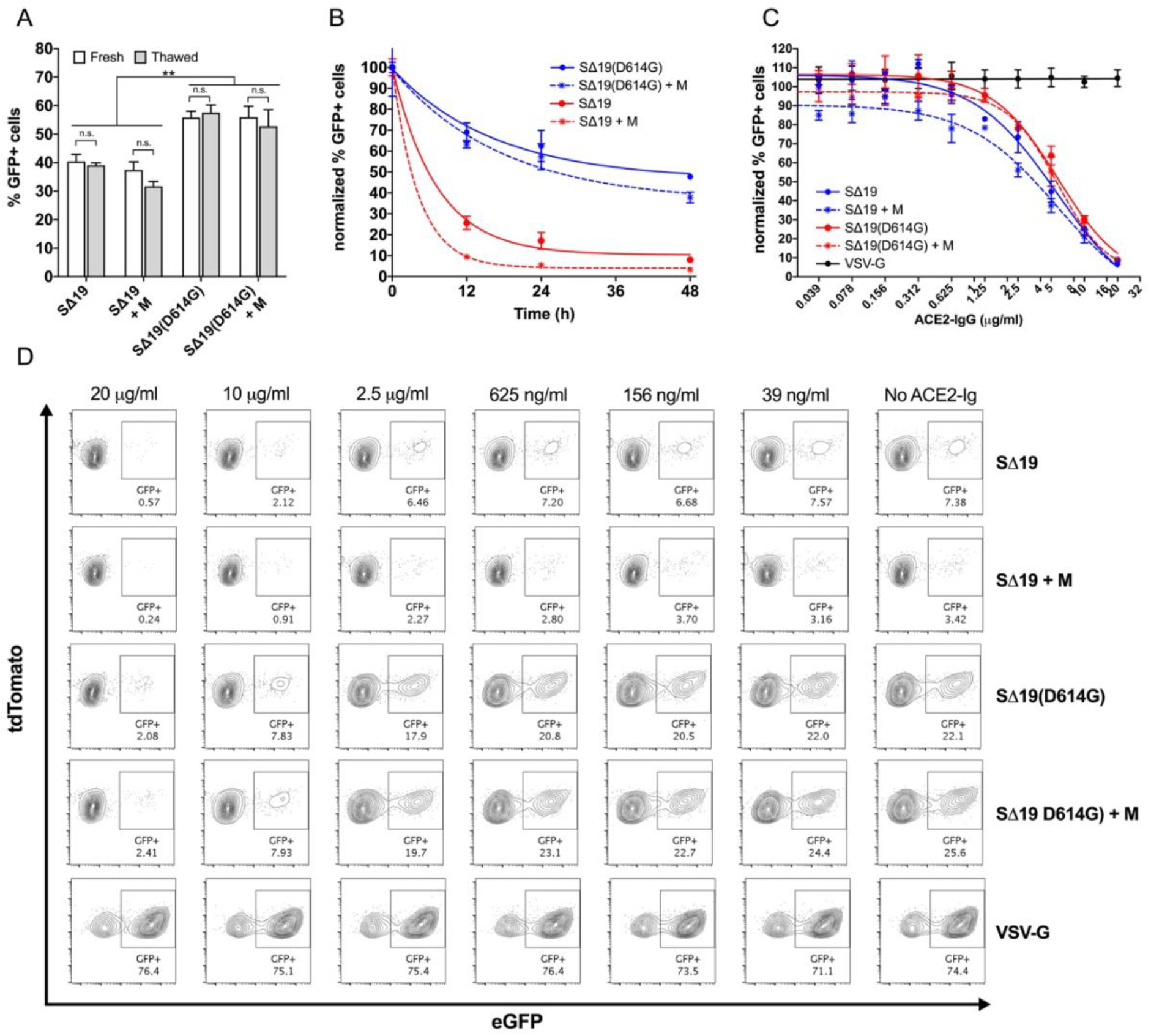
Development of neutralization assay using Spike-pseudoyped lentiviral vectors. **(A)** Percentage of GFP+ 293FT-hACE2 cells after transduction with pseudovirus containing supernatants using the WT or D614G SpikeΔ19 plasmids with or without the M protein plasmid. Transductions are done with either fresh or frozen and thawed viruses. (Results from three different batches of pseudovirus each run in triplicates, ** p<0.005, two-way ANOVA, Sidak’s test) **(B)** Stability of SpikeΔ19 and SpikeΔ19(D614G) pseudoviruses with or without the M protein. Pseudovirus supernatants were incubated at 37°C for up to 48 hours followed by transduction to tdTomato-hACE2 expressing HEK293FT cells. Data normalized to transduction results of freshly thawed supernatant. (Analyzed with one-phase decay model with least squares fit, R2>0.9 for all curves) **(C)** ACE2-IgG mediated neutralization of SpikeΔ19 and SpikeΔ19(D614G) lentiviruses with or without M protein. Spike-pseudotype and VSV-g enveloped lentiviruses were pre-incubated with various concentrations (20μg/ml – 39ng/ml) of ACE2-IgG for 1 hour at 37°C followed by transduction of tdTomato-hACE2 expressing 293FT cells. VSV-G pseudotyped particles incubated with hACE2-IgG were used as control. Graph shows the percentage of GFP expressing cells as normalized to samples transduced without any ACE2-IgG pre-incubation. (Curve fitting was done by 4-parameter non-linear regression using variable slope, R2 values for all curves except VSV-G were above 0.9) (D) Results of flow based analysis of neutralization by ACE2-IgG from one representative experiment.

### Neutralization of pseudovirus particles by soluble ACE2-IgG

Having created four different types of pseudovirus particles, we next sought to determine whether these modifications made any difference in the neutralization of these particles by soluble ACE2-IgG. In order to analyze this, serial dilutions of purified ACE2-IgG were added into the media during the co-incubation of 293FT-hACE2 cells with pseudovirus particles. For each type of pseudovirus, the percentage of GFP+ cells was normalized to the samples where no ACE2-IgG was used. As expected, ACE2-IgG seemed to be specifically blocking the interaction of Spike with hACE2 on the target cell surface since no neutralization effect was observed when using VSV-g pseudotyped particles. No significant difference between the susceptibility of pseudoviral particles to neutralization by soluble ACE2-IgG was observed and the IC50 values calculated using these curves were around 4-8 μg/ml. Comparison of the curves using extra sum-of-squares F test did not reveal any statistically significant difference between the different pseudotypes (**Fig 4C and 4D**).

Taken together, these results show that while the titer and stability of the pseudovirus can change according to the components used, they are still susceptible to neutralization by ACE2-IgG to the same extent.

### Analysis of neutralizing activity in convalescent plasma samples from COVID-19 patients

Having shown the successful neutralization of different pseudovirus particles with soluble ACE2-IgG, we next investigated the neutralizing activity of plasma samples from recovered seropositive COVID-19 patients that volunteered to become convalescent plasma donors. Pseudovirus containing supernatants were co-incubated with donor plasma at various dilutions ranging from 1/20 to 1/20480 at 37°C for 1 hour before they were applied in triplicates to 293FT-hACE2 cells in 96-well plates. Each donor’s plasma was tested against the four different pseudotypes described in this study as well as the VSV-g pseudotype as a control (**Fig. 5A**). Plasma samples from seronegative healthy donors were used as negative controls and showed no neutralization effect on any pseudotype (**Fig. 5B)**.

**Figure 5:**
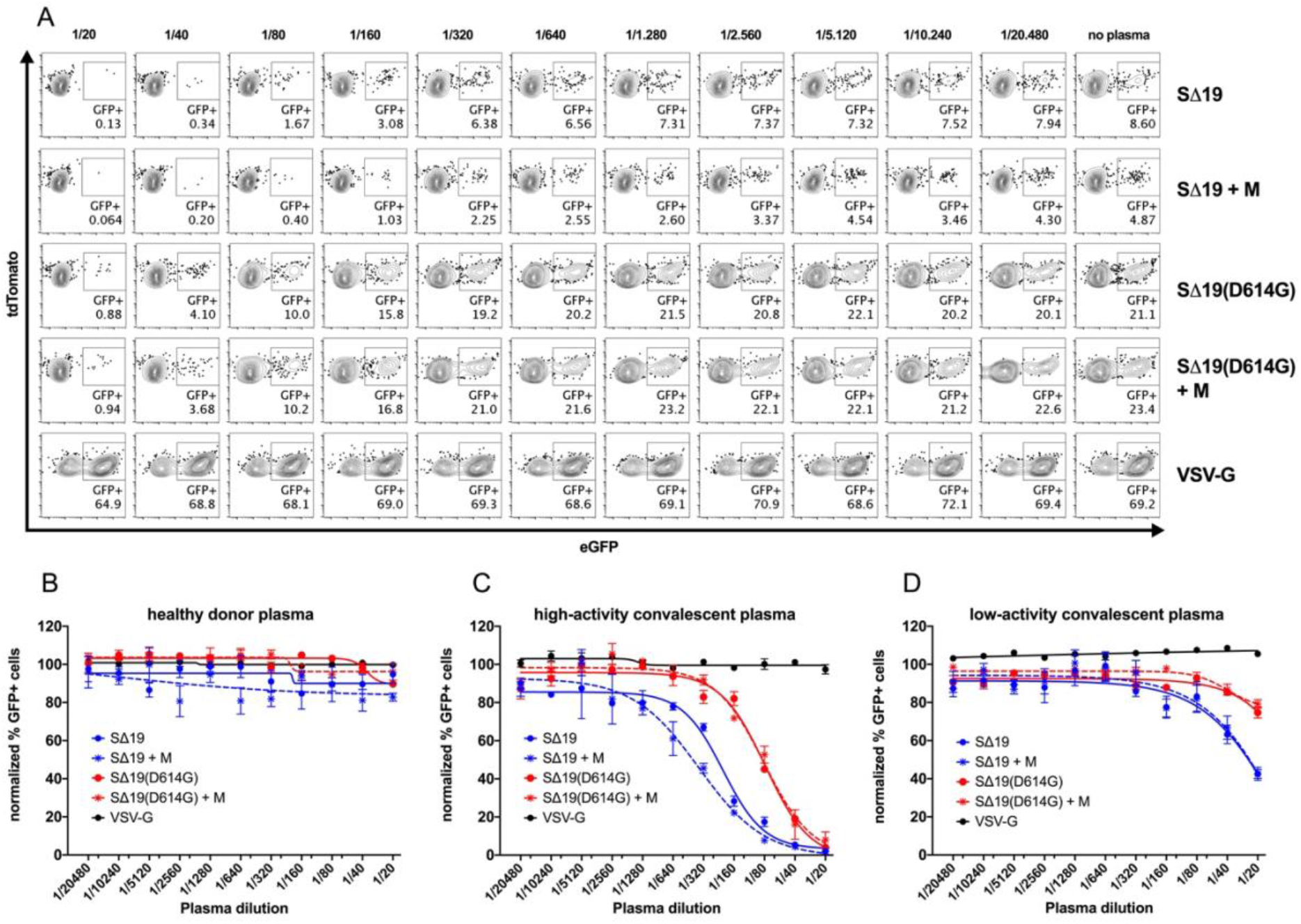
The use of Spike pseudovirus for analysis of neutralizing activity in convalescent plasma samples. Spike or VSV-G pseudotyped particles were incubated with serial dilutions (1:20 – 1:20480) of CP samples for 1 hour at 37°C followed by transduction to 293FT-hACE2 cells. VSV-G enveloped lentiviruses incubated with CP samples were used as negative controls. **(A)** Representative flow cytometry analysis of CP dependent neutralization assay from one donor. Plasma samples from healthy donors and 16 patients that recovered from COVID-19 were analyzed **(B)** Healthy donors are used as a control group and their plasma showed no neutralization effect. (Results from one representative healthy donor) **(C)** 9/16 of CP samples showed high neutralization activity. (Results from one representative high-activity CP) **(D)** 7/16 of CP samples showed no or low neutralization. (Results from one representative low-activity CP) Curve fitting was done by 4-parameter non-linear regression using variable slope.

Using this approach, we analyzed the neutralizing activity of CP samples from 16 donors who had recovered from COVID-19 and were chosen in accordance with the guidelines of the Turkish Ministry of Health. We observed high neutralizing activity in a group of donors (**Fig. 5C**) while some donors showed little-to-no neutralization of the pseudovirus particles (**Fig. 5D)**. Our analysis revealed that 7/16 donors (44%) were in this group with very low neutralizing activity where we were unable to calculate NT50 values due to the fact that even the lowest dilution of plasma used in the assay was unable to neutralize half of the pseudovirus used in the assay. For the remaining 9/16 donors (%56), curve fitting was successful (with R^2^>0.9) and NT50 values were calculated. These results indicate that the extent of neutralizing activity in convalescent plasma samples varies greatly and defining donors by simple IgG positivity might not be the best approach to predict the efficiency of convalescent plasma therapy.

It has previously been reported that the extent of neutralizing activity in convalescent plasma samples correlates with the severity of COVID-19 experienced by the donor, where hospitalized patients presented with much higher neutralizing activity in their plasma samples upon recovery from the disease (51–53). Confirming this observation, we find that none of the donors presenting with low-activity plasma had been hospitalized due to COVID-19 and the extent of symptoms at diagnosis was generally higher in the group with high-activity CP (**Table 1**).

### Neutralizing activity does not necessarily correlate with antibody titers

Besides curve-fitting and calculation of NT50 for the neutralization assays, we further carried out a semi-quantitative analysis of anti-SARS-CoV-2 antibodies in CP samples using WIZ Biotech’s colloidal-gold based rapid detection kit in conjunction with the WIZ-A101 Portable Immunoanalyzer.

Despite observing a variation in the level of IgG and IgM concentrations in all plasma samples, the extent of this positivity did not necessarily correlate with neutralizing activity (**Fig. 6A and 6B**). While some plasma samples with high SARS-CoV-2 specific IgG titers showed high neutralization activity as expected, our results also revealed the presence of donors with high IgG positivity showing no detectable neutralizing activity as well as donors with relatively lower IgG positivity that perform better than expected. This lack of direct correlation suggests that the neutralizing activity of the plasma depends more on the quality of the humoral immune response rather than the quantity of it.

**Figure 6:**
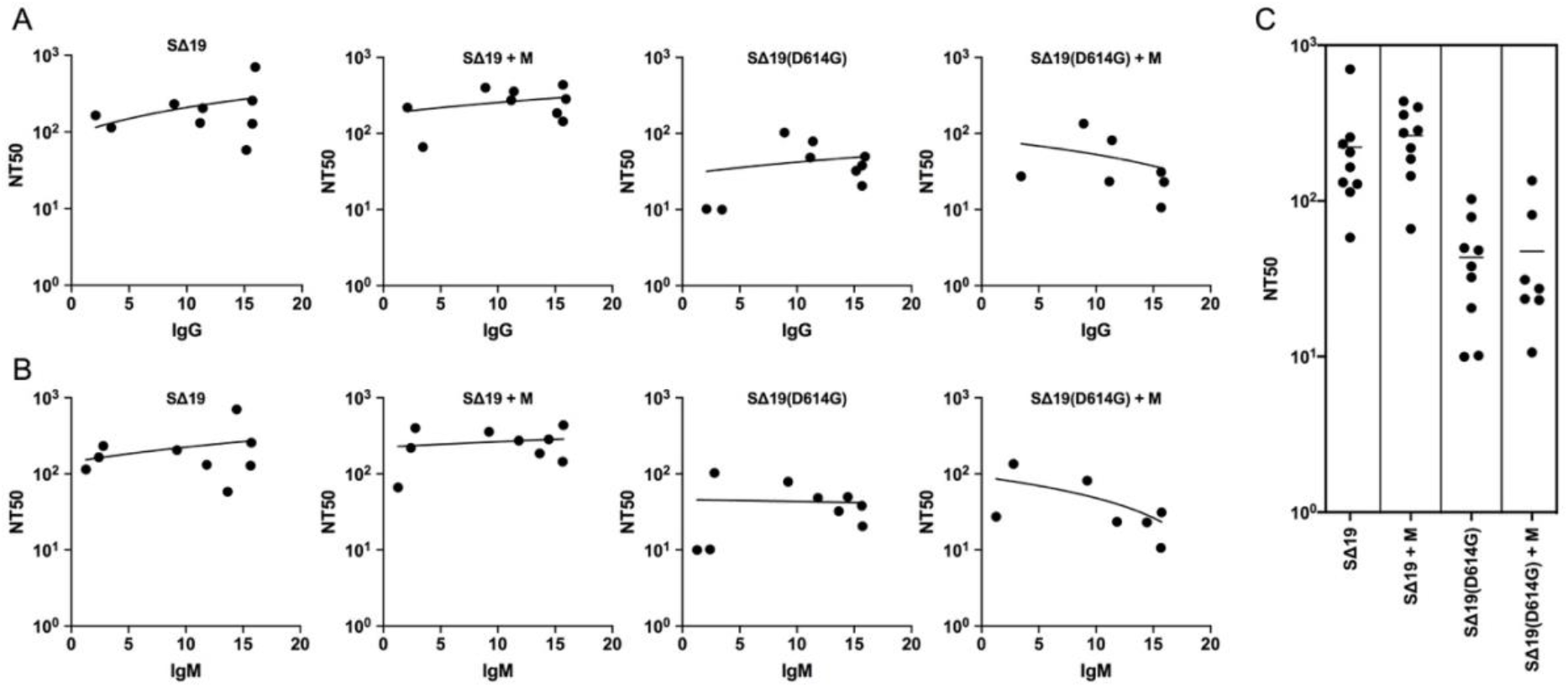
Analysis of neutralizing activity in high-activity convalescent plasma samples. The dilution factor corresponding to the calculated IC50 was used as Neutralizing Titer 50 (NT50) value. **(A)** NT50 values against the various pseudotypes used in this study plotted against IgG positivity measured a semi-quantitative colloidal-gold based rapid detection kit. (Lines show simple linear regression. Deviation from 0: not significant) **(B)** NT50 values against the various pseudotypes used in this study plotted against IgM positivity measured a semi-quantitative colloidal-gold based rapid detection kit. (Lines show simple linear regression. Deviation from 0: not significant) **(C)** NT50 values of high-activity convalescent plasma samples measured against the different pseudotypes used in this study.

Our results also revealed donor-dependent differences in the neutralizing activity of plasma samples against the different virus pseudotypes (**Fig. 6C**). The extent of neutralizing activity against the different pseudotypes did not significantly change but a trend towards lower neutralization of the D614G mutant was observed. While these results are indicative of differential humoral response against the D614G mutant, the lack of knowledge on the genotype of the virus originally infecting the donor prevents us from further speculating on this issue. As a number of studies have already demonstrated increased susceptibility of the D614G mutant to neutralization (27, 54, 55), our observations were regarded as a consequence of the higher titer and stability of the D614G pseudovirus. The response against pseudovirus particles carrying the M protein also varied in a donor-dependent manner and did not have a significant effect on the NT50 values observed.

Taken together, these results indicate that the extent of neutralizing activity in convalescent plasma samples depends on the quality of humoral immune responses and varies greatly between donors. Functional evaluation of neutralizing activity using pseudovirus-based neutralization assays must be accepted as a critical indicator for choosing convalescent plasma donors if clinical efficacy is to be maximized.

## Conclusion

This report presents our efforts to optimize the production of lentiviral vector-based SARS-CoV-2 pseudovirus particles and their use in neutralization assays to measure the activity of convalescent plasma samples. We primarily focused on using different variants of the Spike protein and optimizing transfection and supernatant collection conditions for optimal pseudovirus production. Our results clearly demonstrate that the removal of the ERRS in the tail of the SARS-CoV-2 Spike protein is crucial for efficient lentiviral-vector based pseudovirus packaging and the use of Spike(D614G) mutant significantly increases the titer and stability. We have also demonstrated that inclusion of Sodium Butyrate in the cell culture medium during supernatant collection dramatically increases pseudovirus production.

More importantly, our results indicate that a significant portion of CP donations defined by simple IgG positivity indeed contain very low neutralizing activity. While our sample size was inadequate to analyze clinical responses from the use of such plasma, the extent of neutralizing activity has already been identified as a significant factor in the clinical efficacy of CP therapy (56–58). Despite the imminent arrival of massive COVID-19 vaccination campaigns, a significant level of vaccine hesitancy is unfortunately expected throughout the world (59–61). Coupled with the lack of an efficient drug therapy for COVID-19, this means that treatment modalities such as CP administration or mAb-based therapeutics still need to be perfected and ready-in-place until SARS-CoV-2 is fully eradicated. Pseudovirus-based neutralization assays can play an important role in this process by identifying high-activity CP donations for maximizing clinical benefit.

## Acknowledgements

The authors would like express their gratitude towards Prof. Mehmed Özkan, the President of Boğaziçi University, to Prof. İsmail Cinel, the Chief Physician of Marmara University Pendik Education and Research Hospital and to Prof. Şaban Tekin, the Director of TUBITAK-MAM Genetic Engineering and Biotechnology Institute for their continuing support during the pandemic. The authors would also like to thank Prof. Mayda Gürsel of Middle East Technical University, Prof. İhsan Gürsel of Bilkent University and Prof. Batu Erman of Boğaziçi University for fruitful discussions.

## Author contributions

C.P, E.C., E.Z.E and Z.S.K carried out cloning, cell culture, flow cytometry analysis, pseudovirus production and neutralization assays. G.T. and M.A. carried out pseudovirus production and data analysis. C.E. collected convalescent plasma samples. U.S., H.B. and S.B. contributed to donor recruitment. I.S. carried out western blot analysis and project management. B.C.M. carried out microscopy analysis. B.D., O.T. and G.D.D. carried out purification of recombinant ACE2-IgG. G.T., G.D.D. and N.O. contributed to study design and manuscript writing. T.S. conceptualized the study, analyzed data and wrote the manuscript. All authors reviewed the final version of the manuscript.

